# Congruent timing in Human, Nightingale, Parotid Wasp and Hawaiian Planthopper vocalisations

**DOI:** 10.1101/2024.07.15.603552

**Authors:** Adrian Fourcin

## Abstract

Earlier multi-speaker, multi-language measurements found that in purposeful fluent speech, total voice duration was half that of the whole – a maximum Shannon entropy criterion for two state signals. The present aim is to discover whether other species’ vocalisations are similarly associated with maximum timing entropy control. Data selection has been based on the use of courtship songs since they are purposeful and fluent. These songs are intended for con-specific females and, at present, often too complex for reliable acoustic signal analysis. In consequence it has been necessary, for this preliminary study, to find courtship songs in which the echemes are well demarcated from silence – this has led to a curtailment of potential signal resources.

The vocalisation timings in examples of the courtship songs of Nightingale, Parotid Wasp and Hawaiian Planthopper have been analysed. The resulting measurements are compared with those for voice timing in purposeful fluent, human speech. There are three primary findings:

- The same Shannon timing entropy maximising rule is intrinsic to all four vocalisations.
- Gaussian distributions of voice/vocalisation timings are present for each of the four signal types.
- In each case, maximum entropy of voice/vocalisation timing tends to be maintained continually throughout the production of the whole signal.

## Introduction

This study of cross species voice timing has its origin in work to extend a set of speech signal analytic tools for pathological voice measurement^1^. (*“Voice” is associated, here, with the pulsatile pressure excitation of the speaker’s vocal tract produced by the interruption of the pulmonic air stream by vibratory closures of the laryngeal vocal folds*.) Acoustic speech pressure and simultaneously acquired electrolaryngograph vocal fold contact waveforms^1^ were used to provide physical correlates of the range and regularity of patients’ loudness, pitch, voice quality and, more recently, fluency.

Fluency was assessed on the basis of the temporal measurement in connected speech of the relative durations of: voice in sonorants and voiced obstruents^1,2^ (V), relative to the combined durations of voiceless obstruents^1,2^ (A), and silence (S).

It was noted informally that achieving an equality of timing between the sums of (V) and (A+S) was one aspect of a return to speech production normality. In Shannon’s mathematical theory of information^3^ this temporal equality in a two state signal source is an indication of maximum entropy. A controlled set of measurements was made to investigate whether this simple timing rule was well established in normal fluent purposeful speech^4^.The resulting analyses of recordings made by 56 dramatic arts students and academic speakers from eight different language backgrounds gave strong evidence for the existence of an intrinsic tendency to achieve maximum voice timing entropy. These original voice timing measurements were based on the use of vocal fold closure detection signals obtained and analysed with Laryngograph^®,1^ hardware and software in terms of echeme timings^7^. A confirmatory model Matlab script^5^ has been written for the present work.

The present exploratory research aims to discover if there is quantitative evidence that the precise voice timing control rule found in purposeful fluent human speech^4^ might also be found in the fluent meaningful vocalisations of other species. Courtship song in other species may provide a source of data that involves crucially important meaningful fluent vocalisations (echemes^7^). Three examples have been analysed, in terms of echeme timings, making use of the model Matlab script^5^, with individual adjustments for sampling frequency, signal to noise ratio and data source.

Temporal structures of an early morning recording of the courtship song of the male nightingale (recorded by Nick Penny^8^) have been measured. This song is typically produced as a sequence of vocalisations, each differing from the others in duration and complexity and followed by silent intervals of different durations. This temporal vocalisation organisation is similar to that of human voicing (V) in the prosodic sequences of connected speech, with the difference, as noted above, that human speech also makes structured use of voiceless obstruent sounds (A). Measurements based on the model speech script^9,10^ have been made of this nightingale data. In the same way as for all the reported speech data, a close approximation to the ideally encoded 50% ratio of vocalisation timing to total song duration was found.

A recording of the courtship song of the male Parotid wasp has been published^11,12^, and its acquisition discussed with the authors. Successive wing flaps have been controlled by the wasp to produce acoustic pulses of varying complexity and duration (echemes^7^), separated by varying durations of silence. This basic temporal structure is similar both to that in human voice and vocalisations in the courtship song of the nightingale. A script^13^ based on the model used for voice analysis^5^ has been applied to measure the relative probabilities of vocalisation and silence in the whole recording. A close approximation to the ideally encoded 50% ratio of vocalisation timing to total song duration was found.

A description of the courtship song of the Hawaiian Planthopper has been published^15^ and its acquisition discussed with its principal author who has pointed to the publicly available archive containing the field recordings^16^. An extension of the exemplar script^18^. has been used to measure the relative probabilities of vocalisations (echemes) and silence in this complete courtship song^18^. Once more, there was a close approximation to the ideally encoded 50% ratio of total vocalisation timing to the total song duration.

Further analyses, and extensions, of all of these echeme timing measurements have found that both full and part-data recordings are approximately Gaussian distributed.

## Materials and methods

### Speech based data and scripts

The original voice timing measurements, on which this work is based, employed vocal fold closure detection signals obtained and analysed with Laryngograph^®,1.4^ hardware and software. A model Matlab script^5^ has been written for the present work and applied to verify the original voice timing measurements using the recorded acoustic speech and laryngograph signals from one of the original speech data files^4,^ see discussion in Figure 1. All the previous speech based measurements^4^. were based on the use of normal fluent purposeful speech. Courtship songs from other species have been used to provide comparable fluent, purposeful, vocalisation data recordings. Three examples^9,12,17^ have been analysed in terms of echeme^7^ timing, making use of scripts^10,13,18^ each based on a model human voice analysis script^5^, with individual adjustments for sampling frequency, signal to noise ratio and data source.

**Figure 1.**
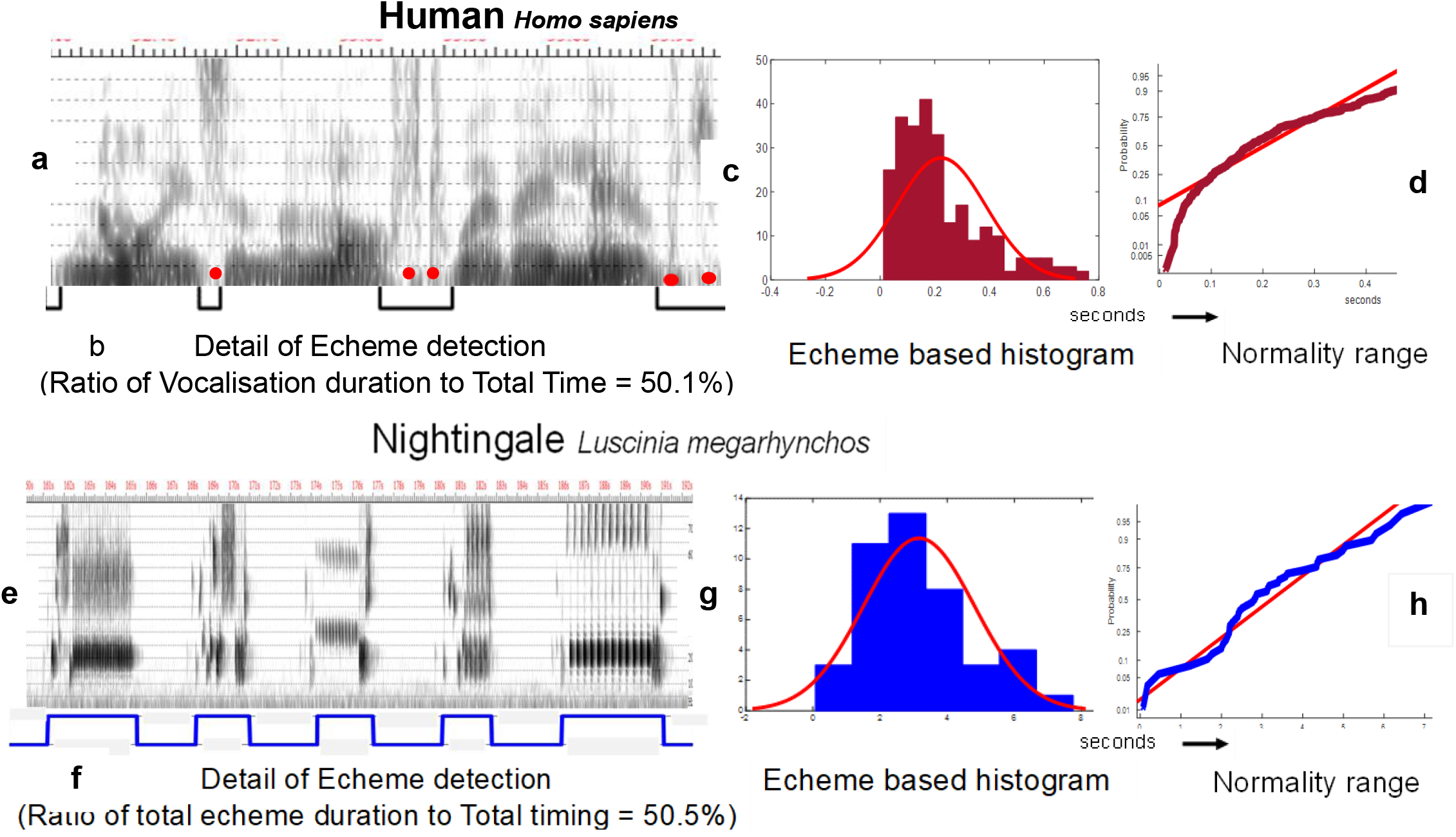
Human Voice and Nightingale Courtship Song signal analyses

### Nightingale data and scripts

An early morning recording of the courtship song of the male nightingale, recorded by Nick Penny^8^ has been included with his permission. The temporal structure of echemes in this data^9^ has been measured using an adjusted voice analysis model script^10^

### Parotid Wasp data and scripts

A recording of the courtship song of the male Parotid Wasp has been published^11^, and its acquisition discussed with the authors. The echeme timing I this data^12^ has been analysed using an adjusted script based on the model used for voice analysis^13^

### Hawaiian Planthopper data and scripts

A description of the courtship song of the Hawaiian Planthopper has been published^15^ and its acquisition discussed with its principal author who has pointed to the publicly available archive containing the field recordings^16,17^. An adjusted extension of the model script^18^. has been used to measure, in the same way as for the other courtship songs, the relative probabilities of vocalisation (echemes) and silence in this complete courtship song.

### Data and Script file structures

In all four sets of data^6,9,12,17^, and for all four scripts^5,10,13,18^. **“1”** at the end of a file name indicates a full original recording, whilst **“2”** indicates that only the first part of the full file is included.

The 4kHz broad band 300Hz spectrogram of Figure 1a is based on the analysis of a 1.9s speech acoustic sample, (“noise in the loft where they lived”) taken, with the speaker’s permission, from a combined speech acoustic and electrolaryngograph^®1^ purposeful fluent recording of 113s duration. The black pulse train in Figure 1b gives an example of the laryngograph^1^ (vocal fold vibratory contact) based, positive pulse, identification of individual voice echemes^5^. This has been used to measure the voice to total speaking time ratio for the whole recording. The total timing includes silences together with aperiodic/voiceless sounds, shown by red dots, of which **[s], [ft]** and /v/ and /d/, here realised as **[ft]**, are particular examples. In the original measurements^4^ Laryngograph^®^ software was used to determine the voice to speaking time ratio of ∼50%. A confirmatory model script has been used here to check the original analyses with reference constants set to adjust for the original sampling frequency and recording signal to noise ratio^5^. The histogram of Figure 1c is based on the analysis of the 257 echemes identified in the whole recording by the analysis script^5^. The mean echeme duration of 0.223s is visually correlated with the mean of the overlaid normal distribution. Although there is no obvious Gaussian shape similarity, The Kolmogorov Smirnov (KS) test^5^, however, does indicate a Gaussian structure and this is visually supported by the partial adherence of the Matlab data probplot to the ideal red normality line in Figure 1d.

**Figures 1 e to h** are based on the analysis of a complete open source male nightingale dawn courtship song (recorded by Nick Penny^9^), total duration of 269 seconds with 43 echemes. Fig. 1e shows an 8kHz broad band 300Hz spectrogram of a 32s, five echeme excerpt whilst Fig 2b illustrates the process of automatically identifying individual echemes using an adjusted model signal processing script^10^. In earlier speech based measurements^4^ maximum timing entropy was achieved by the speaker’s use of laryngeal voicing control that adjusted total voice time to be half that of the total speaking time duration. The ratio of the total duration of all echemes in this song to the song’s total duration is 50.5%. This indicates that, for this particular example, there is a close correspondence between this aspect of nightingale vocalisation and the human voice timing control previously measured^4^, and the particular example of figure 1b above. The histogram of Figure 1g has a mean echeme duration of 3.16s, a value that could be derived from a range of echeme distribution types. The overlaid normality distribution appears, however, to be a good fit to the data. and this is confirmed by a positive KS test^10^. The Matlab probplot of figure 1h, in comparison with the red reference normality distribution line, gives a more convincing indication of the likelihood that this nightingale data has a Gaussian distribution, a criterion for maximally efficient, entropy based, encoding^19^.

**Figure 2.**
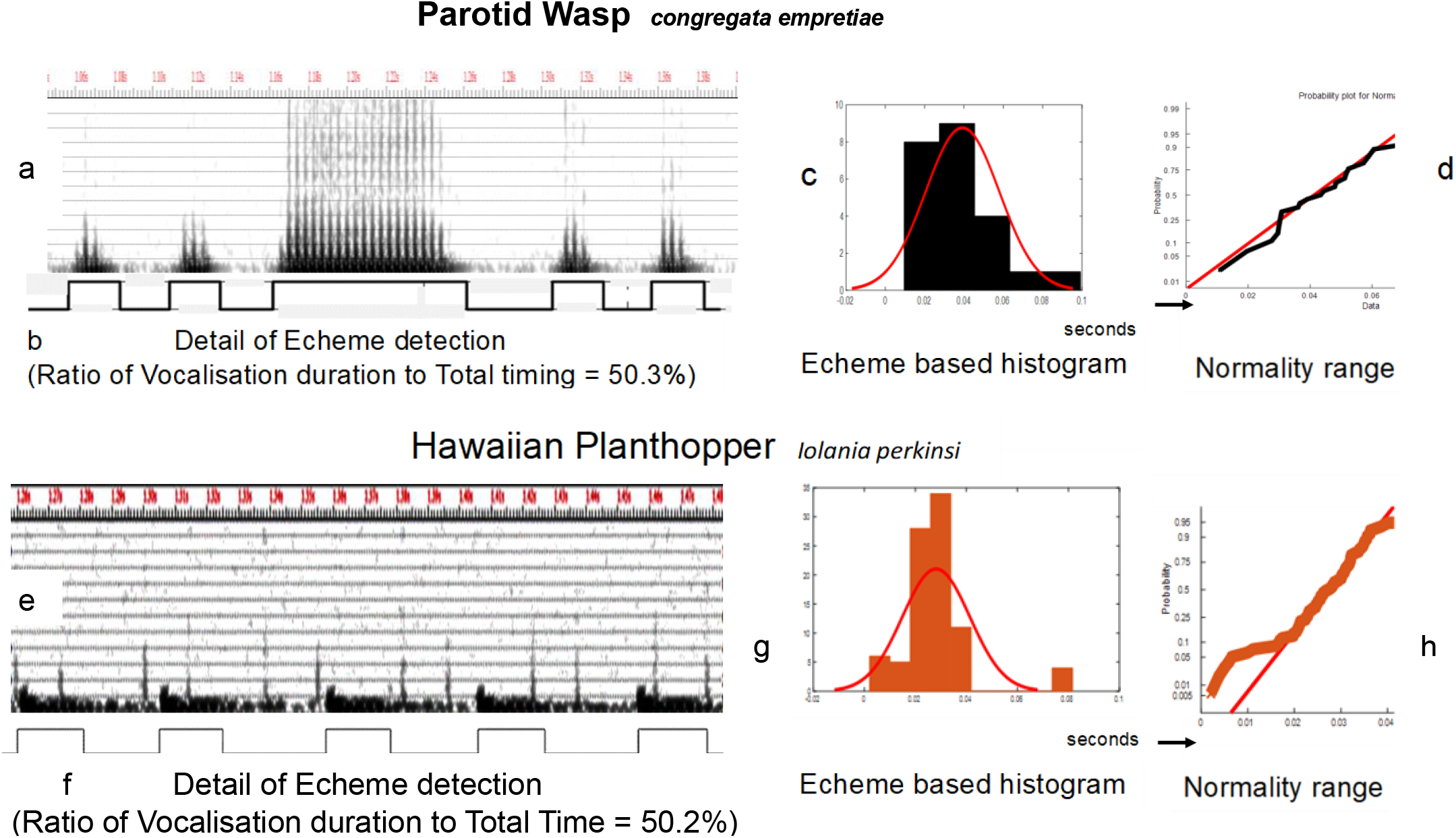
Parotid Wasp and Planthopper courtship song analyses

Figures 2 a to d are based on openly available data acquired in the field by Justin Bredlau^11,12^. The analysed song is the second of a group of three in the original recording. The first and last of the group had malformed coalescing wing flap pulses and are not reported. In Figure 2a the 12kHz broad band 300 Hz spectrogram of 0.34s duration has been taken from a total signal duration of 1.79s and shows five echemes from a total of 23. These spectrogram linked echemes correspond to the positive peaks of the waveform in Figure 2b following their simple amplitude level detection by the Wasp version of the model script^13^.

Manual analysis is also feasible using the quite separate WASP program^14^, but the model script, adjusted for the parameters of this recording, gives ready access to the histogram analysis of Figure 2c and the mean wing flap duration measurement of 0.039s. This mean duration corresponds, once more, to a total echeme duration ratio of 50.3% of the total duration of the whole song^10^. The overlaid red plot gives an indication of normality compliance. This is more accurately confirmed by a KS analysis and the close Gaussian approximation of this data sample to the ideal normality reference red line shown in Figure 2d

Figures 2 e to h are based on the analysis of a brief isolated complete Planthopper courtship song of 4.9s duration extracted from field recordings^17^ by Hannelore Hoch – who reported that courtship songs can be hours long. The 22kHz 300Hz analysing bandwidth broadband spectrogram of Figure 2e shows five echemes of a total of 88. The waveform peaks of Figure 2f are based, as in all the other figures, on the use of the original more clearly defined acoustic waveform^18^. The average echeme duration is 0.028s and the total duration of all echemes is 2.47s again corresponding to a total echeme to overall duration of 50.5%. Both the KS test and the overlaid normal distribution in Figure 2g indicate that the distribution of echeme durations is largely Gaussian compliant and this is visually confirmed by the close adherence of the data plots in Figure 2h to the ideal normality red reference line. In addition to the data for each of the songs in these four figures, data for half way recordings has also been placed in the data repository ^6^,^9^,^12,17^. For all four of these half way data recordings there is a close correspondence between the ideal normal distribution lines and the Matlab probplots for normality range (see Figure 3). These measurements indicate that each signal source exerts continuous monitoring throughout the whole process of vocalisation production of the echeme durations so as, consistently, to approximate maximum entropy. encoding in real-time without transmission delay.

**Figure 3.**
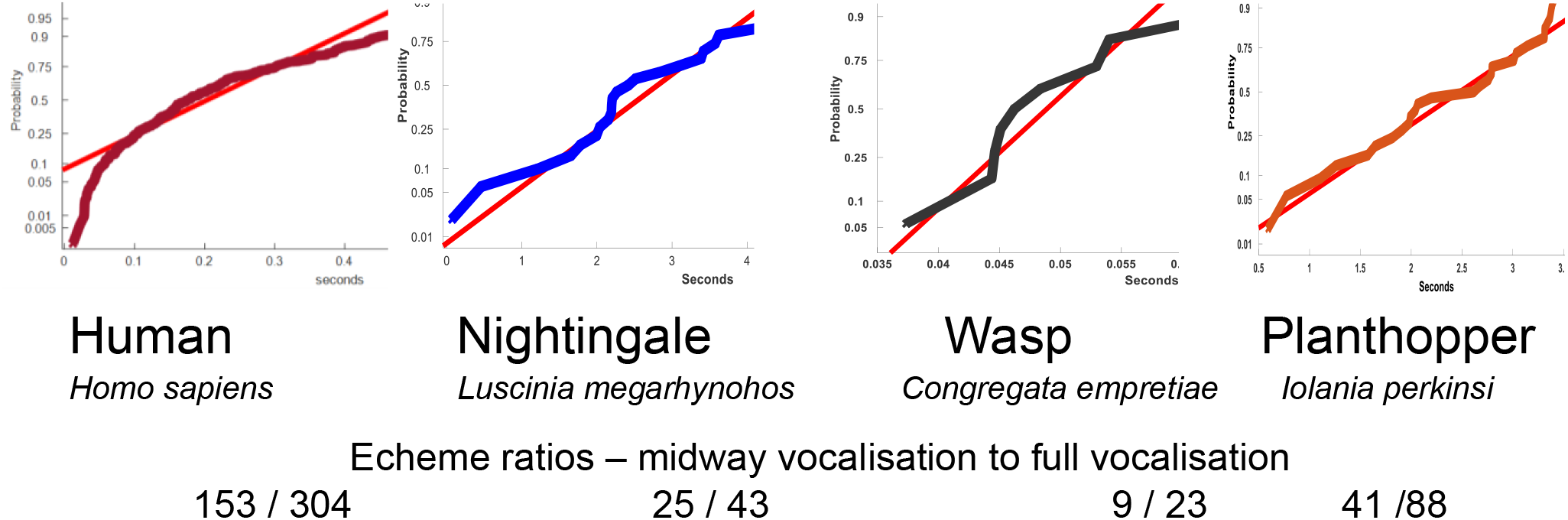
Mid Way Normality Ranges

## Discussion

This work follows a chance observation that in human purposeful fluent speech, voice interval durations were organised so as to comply with a simple Shannon entropy maximising rule.

The original aim of the present study was to examine whether other than human species, in purposeful fluent communication, controlled the timing of their vocalisations to comply with a V = S rule, where V is the total duration of vocalisation and S is the total duration of silence in the whole utterance. Vocalisation timing in particular examples of Nightingale, Parotid Wasp and Hawaiian Planthopper (N, W, P) courtship songs were analysed, since these songs are likely to be both fluent and purposeful. Close approximations to V = S equality have indeed been found, corresponding to the V = (A + S) rule that was found in more extensive measurements with human purposeful fluent speech^4^ – where A is the total time taken by aperiodic, voiceless, speech sounds.

Closer examination of all the data found that all of the measured vocalisation timing distributions were quite closely Gaussian, irrespective of the mechanisms of vocalisation. This is a requirement for maximum signal entropy^19^, and the measured timing equalities are a reflection of that compliance.

Efficient signal transmission in commercial practice frequently makes use of entropic encoding^20^. This involves both processing the signal’s information stream and the introduction of concomitant delay between transmission and reception. The ecological examples of voice/vocalisation based communication discussed here are not, however, subject to this encoding disadvantage. In each case, the signal source is able to control the timing of its information stream so that efficient encoding is feasible in real time. Particular examples are included in the associated “2” format data and script files^6,9,12,17^ ^&^ ^5,10,13,18^ of the mid-stream analyses of each of the human and NWP data files. Figure 3 shows their corresponding Matlab probplot normality ranges. The full file versions are closely comparable – see Figures 1 and 2.

A finding of special interest is that timing in human voice production appears also to be controlled in real-time to achieve maximum voice timing entropy. This implies that both sonorants^1^ and voiced obstruents^1^ have a special place in the evolution of human acoustic communication. The complementary optimal temporal organisation of voiceless obstruents^1^ remains, however, to be confirmed.

A major weakness of the work comes from its use of a very small data set. The quantitative measurements, however, appear compelling. An examination of their more general cross species validity may provide a basis for more extended research, using resources that have not been available here. Since efficient communication can provide vital support for survival, the same tendency to optimise the statistical probabilistic structures of communication in other species, and for other than durational data, may be found more generally. The chordata, hymentoptera and arthropoda phyla diverged ∼700MYA^21^ and it might be supposed that these strategies of optimal signal encoding originate with ancestors of even greater antiquity. However that may be, it seems possible that the present observations stem from the operation of a well-defined intrinsic signal processing structure that may be found in other species and genera.

